# Chronic wireless streaming of invasive neural recordings at home for circuit discovery and adaptive stimulation

**DOI:** 10.1101/2020.02.13.948349

**Authors:** Ro’ee Gilron, Simon Little, Randy Perrone, Robert Wilt, Coralie de Hemptinne, Maria S. Yaroshinsky, Caroline A. Racine, Sarah Wang, Jill L. Ostrem, Paul S. Larson, Doris D. Wang, Nick B. Galifianakis, Ian Bledsoe, Marta San Luciano, Heather E. Dawes, Gregory A. Worrell, Vaclav Kremen, David Borton, Timothy Denison, Philip A. Starr

## Abstract

Invasive neural recording in humans shows promise for understanding the circuit basis of brain disorders. Most recordings have been done for short durations from externalized brain leads in hospital settings, or from first-generation implantable sensing devices that offer only intermittent brief streaming of time series data. Here we report the first human use of an implantable neural interface for wireless multichannel streaming of field potentials over long periods, with and without simultaneous therapeutic neurostimulation, untethered to receiving devices. Four Parkinson’s disease patients streamed bilateral 4-channel motor cortical and basal ganglia field potentials at home for over 500 hours, paired with wearable monitors that behaviorally categorize states of inadequate or excessive movement. Motor state during normal home activities was efficiently decoded using either supervised learning or unsupervised clustering algorithms. This platform supports adaptive deep brain stimulation, and may be widely applicable to brain disorders treatable by invasive neuromodulation.

## Introduction

Electrical stimulation using permanently implanted brain devices has become a standard therapy in movement disorders and epilepsy, and is also under active investigation for psychiatric and cognitive disorders^1^. Often, neurostimulation therapies are introduced without a clear model of the underlying circuit disorder, nor of the mechanism by which therapeutic stimulation influences signs and symptoms. One approach to addressing this knowledge gap has been analysis of invasive cortical or subcortical recordings from externalized leads, either during lead implantation surgery or for a few days afterwards in the hospital setting. This method provides neural data with excellent signal to noise ratio at high spatial and temporal resolution compared to noninvasive methods^2^, but is limited by its short duration, unnatural environment, and temporary circuit changes induced by edema from recent surgery. There is thus interest in incorporating a sensing function into chronic, fully implanted neurostimulators, for chronic neural recording over months or years^3^. In addition to circuit discovery, an exciting potential applications for these “bidirectional” (sense and stimulate) neural interfaces is adaptive neurostimulation, in which stimulation therapy is automatically adjusted in response to changing brain states, decoded from electrophysiological biomarkers of symptom severity^4,5^.

However, the early generation bidirectional neural interfaces have important limitations^4-6^. Bandwidth and duration of data collection for time series data have been limited. Wireless data streaming typically requires patients to be “tethered” to a receiving interface that restricts free movement, and requires the presence of trained investigators, constraining the use of such devices to unnatural environments. Large stimulation artifacts and amplifier saturation may preclude data collection during therapeutic stimulation. Here we report the first human experience with an investigational second generation bidirectional interface, Summit RC+S (Medtronic), which solves many of these limitations^7,8^. This device can transmit neural data at a sampling rate up to 1000 Hz to an external Windows-based tablet up to 12 meters away, allowing freedom of movement. It can be tailored for easy home use and for different disease indications, by programming customized functions within its application programming interface. The device’s recharging capability obviates concerns inherent in prior devices that extensive sensing would lead to premature battery failure. Chronic wireless data streaming from RC+S has only been tested in canine^8^ and nonhuman primate models^9^ of epilepsy. This and similar devices now under development allow researchers and clinicians access to large “real world” neural data sets to enable discovery of personalized neural signatures of human brain disorders.

Patients with Parkinson’s disease (PD) are an ideal population for testing novel bidirectional interfaces. PD is commonly treated with deep brain stimulation (DBS) of basal ganglia nuclei to treat motor fluctuations, the tendency of patients to cycle between bradykinetic (slow), normal, and dyskinetic (excessive movement) states, in response to dopaminergic agents^1^. Short term testing using externalized leads has suggested that adaptive DBS could offer improved efficacy^10^ and reduced adverse effects^11^ compared with standard, continuous DBS. There is a candidate biomarker for bradykinesia identified from acute, peri-operative recordings from externalized leads: the amplitude of beta band (13-30 Hz) oscillatory activity in basal ganglia field potential recordings^2^. However, cortical biomarkers of parkinsonian motor signs have also been proposed^12-14^ and it is not known which site (cortical or basal ganglia) is most effective for motor state decoding. It is unclear whether biomarkers identified in hospital settings remain useful in naturalistic environments, on patients’ regular schedule of dopaminergic medications, or over longer periods.

To address these questions, we implanted four patients bilaterally with RC+S devices attached to both subthalamic nucleus (STN) and motor cortical leads (**Figure 1**). Patients streamed simultaneous multisite field potential data for many hours at home, paired with wearable monitors for independent validation of motor state, with and without therapeutic DBS. We utilized analytic methods that leverage high volume data sets, including both supervised and unsupervised clustering methods, to refine and extend the “oscillation model” of movement disorders pathophysiology^2^. We demonstrate for the first time that oscillatory phenomena in PD correlate with individual patients’ motor signs at home during everyday activities, and that multiple recording sites improve the classification of patient motor state. Using PD as a model, we illustrate capabilities of this novel class of bidirectional neural interfaces, that may be widely applicable for circuit discovery and rational design of neuromodulation therapies in neurology and psychiatry^15^.

**Figure 1.**
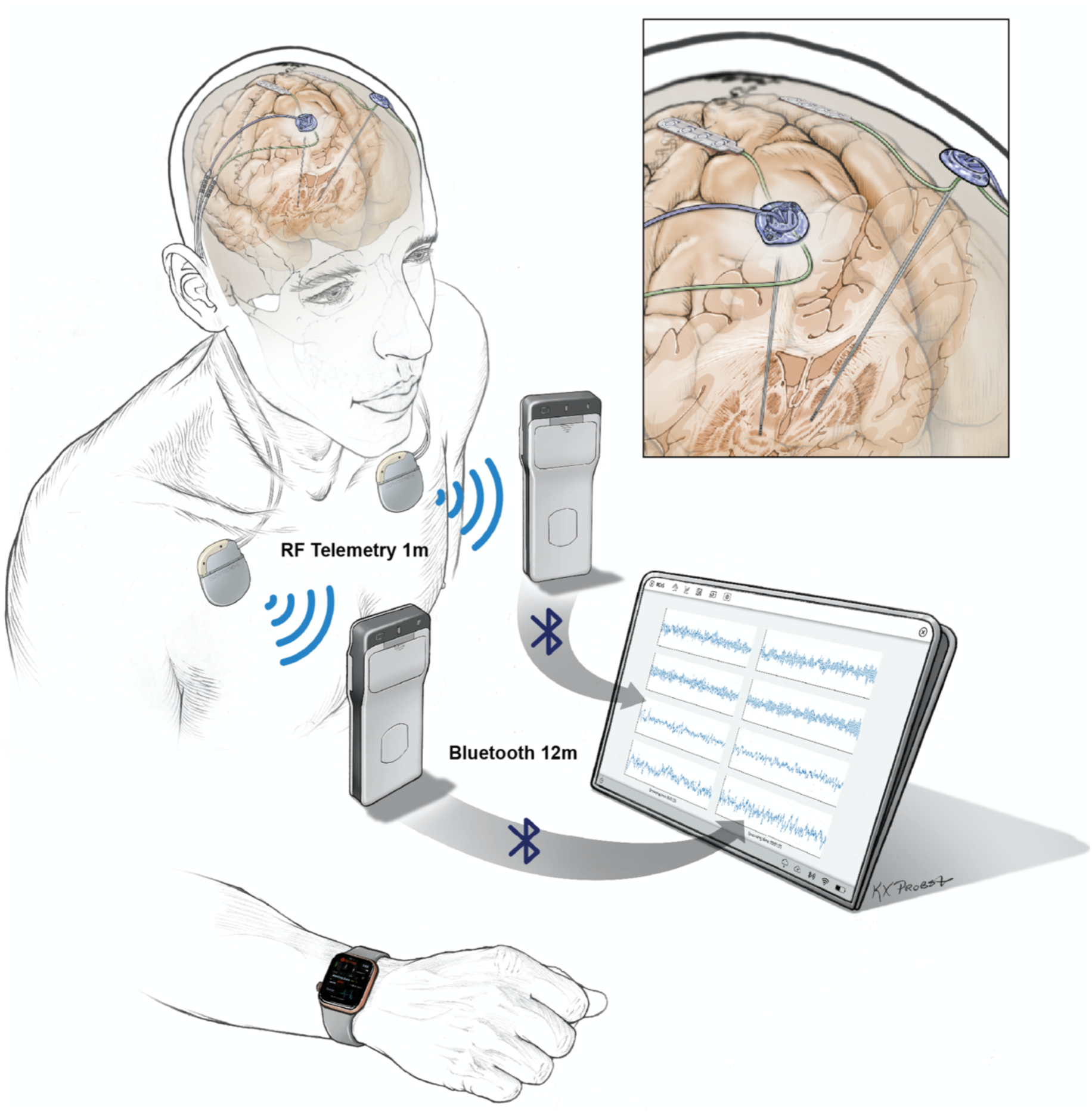
Configuration of implanted hardware and method of data streaming. Quadripolar leads were placed bilaterally into the subthalamic nuclei and in the subdural space over precentral gyri (inset provides zoomed-in view). Each DBS lead and cortical paddle pair were connected via tunneled lead extenders to the ipsilateral Summit RC+S bidirectional implantable pulse generator (IPG), placed in a pocket over the pectoralis muscle. Each RC+S uses radiofrequency telemetry in the medical implant communication spectrum (MICS) band to wirelessly communicate with a pocket sized relay device, usually worn on the patient. The relay devices transmit by Bluetooth to a single small Windows-based tablet at a distance of up to 12 m, allowing sensing of local field potentials from up to four bipolar electrode pairs for up to 30 hours per IPG, before recharge is needed. Custom software on the tablet allows remote updating of device streaming parameters or adjustment of embedded adaptive DBS algorithms, at home. A wristwatch-style actigraphy monitor is downloaded to a server that is synchronized off-line with neural recordings for brain-behavior correlations.

## Results

### Patient characteristics, surgical implant, and contact localization

Four adults received bilateral implants of the Summit RC+S bidirectional neural interface, attached to quadripolar depth leads in the subthalamic nuclei and subdural paddle-type leads over primary motor and sensory cortices. All study subjects had idiopathic Parkinson’s disease with motor fluctuations, including prominent bradykinesia and rigidity in the off-medication states, but varied with respect to the degree of off-period tremor and on-period dyskinesia (**Table 1**). Data were collected in the first 1-3 months after surgery, by wireless streaming of multichannel field potentials to an external computer, both at home and in-clinic (**Figure 1**).

**Table 1.** Demographics of five implanted subjects under NIH UH3 “Closed loop deep brain stimulation in Parkinson’s disease”. All clinical rating scores are obtained preoperatively within 90 days of DBS implantation.

| Subject ID | Age at surgery and gender | Duration of motor sx (years) | MOC A | levodopa equivalents* | UPDRS III off medication score | % change UPDRS when on | time with dyskinesia** | off tremor scores*** |
| --- | --- | --- | --- | --- | --- | --- | --- | --- |
| RCS01 | 54,M | 7 | 26 | 1425 | 49 | 90% | 4 | left:2<br>right:0 |
| RCS02 | 63,M | 19 | 30 | 955 | 45 | 51% | 1 | left: 3<br>right: 4 |
| RCS03<br>**** | 28,F | 12 | 27 | 1550 | 61 | 73% | 2 | left: 12<br>right: 9 |
| RCS04 | 40,M | 4 | 30 | 1314 | 41 | 65% | 1 | left: 6<br>right: 3 |

\*levodopa equivalents calculated with reference to:
Tomlinson CL1, Stowe R, Patel S, Rick C, Gray R, Clarke CE, Mov Disord. 2010 Nov 15;25(15):2649-53. doi: 10.1002/mds.23429. Systematic review of levodopa dose equivalency reporting in Parkinson's disease.
\*\* UPDRS part IV item 4.1
\*\*\* sum of all updrs-III tremor scores for each side, off medication (UPDRS-III items 16a through 17d)
\*\*\*\*positive for Parkin mutation
sx - symptoms
MOCA - Montreal Cognitive Assessment

Accurate lead placement was verified both by intraoperative physiological recordings (**Figure 2 a,b**) and anatomically by postoperative CT scan computationally fused with preoperative MRI scan (example lead locations **Figure 2c;** lead locations for all patients are provided in **Table S1)**. Therapeutic continuous STN neurostimulation (standard clinical therapy) was initiated at one month post-implantation. There were no serious adverse events related to surgery or to the study protocol.

**Figure 2.**
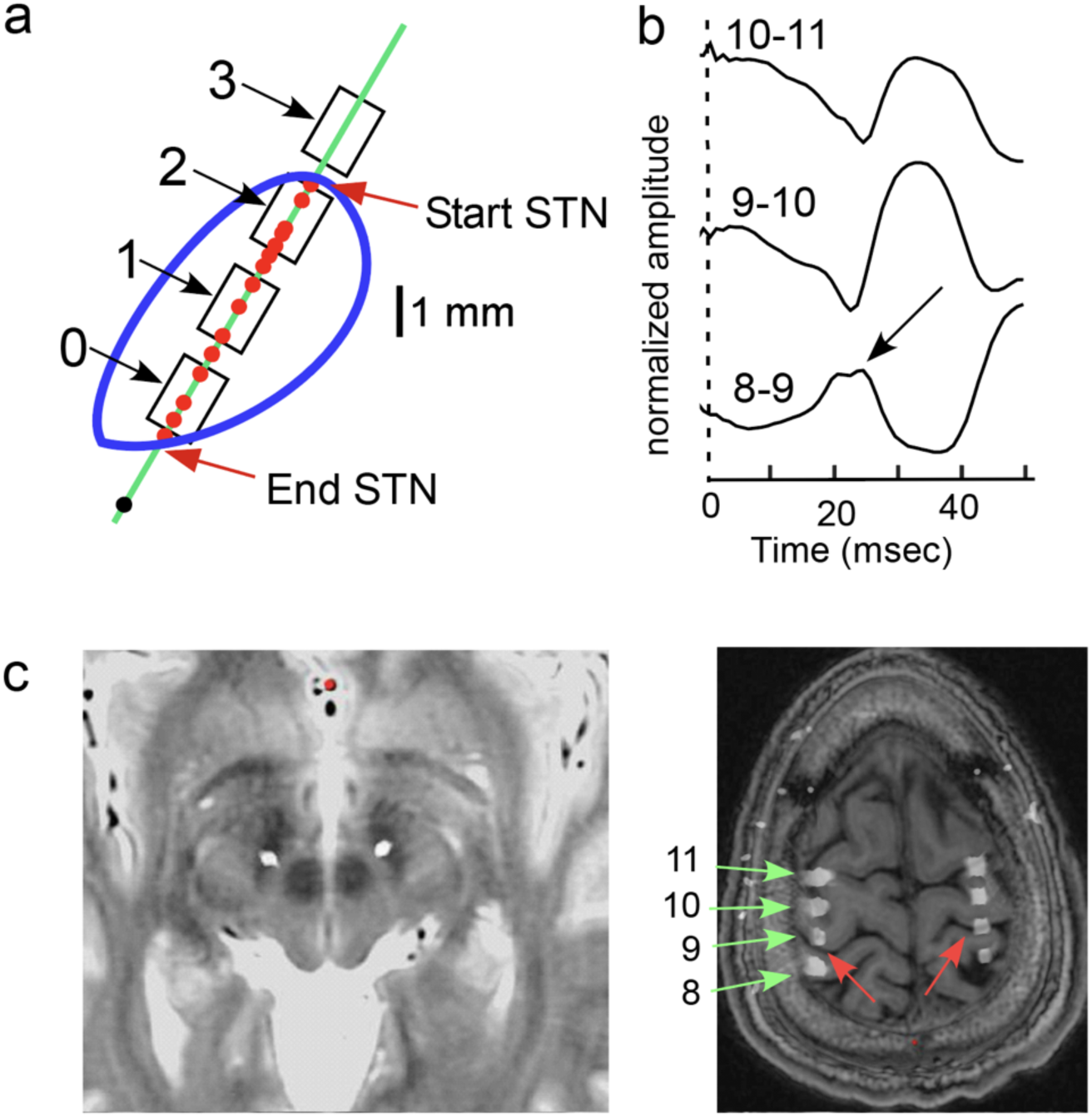
Anatomic and physiological localization of subthalamic and cortical leads. (example from RCS04). **a**, Localization of STN contacts with respect to the borders of STN (outlined in blue) as defined by microelectrode mapping. The microelectrode map (green line) shows the borders of STN as defined by cells (red dots) that have canonical STN single unit discharge patterns and rates. The intended depth of the DBS lead is determined by this map, and contact numbers are labelled. The middle contacts (1 and 2) are within the dorsal 4 mm of STN (motor territory). The black dot is a cell in substantia nigra, pars reticulata. **b**, Somatosensory evoked potential (from stimulation of the median nerve) recorded from the subdural paddle lead, montaged for three overlapping contact pairs. Reversal of the N20 potential between pairs 8-9 and 9-10 (arrow) shows localization of contact 9 to primary motor cortex, consistent with subsequent imaging. **c**, Location of the leads from postoperative CT computationally fused with the preoperative planning MRI. Left, STN leads on axial T2 weighted MRI which shows the STN as a region of T2 hypointensity. Right, quadripolar subdural paddle contacts on axial T1 weighted MRI showing relationship to central sulcus (red arrows) and numbering of contacts (green arrows).

### Data characteristics, movement related activity, and effect of levodopa in clinic

Data were streamed in clinic at a three weeks after implantation, to verify recording quality, the presence of movement related activity, and the effects of levodopa in defined on/off states. Four-channel time series recordings, two cortical and two subthalamic, were done on each side (**Figure 3a**). In sensorimotor cortex, initiation of movement was associated with canonical movement-related reduction in beta band activity with a concomitant increase in broadband activity 50-200 Hz reflecting local cortical activation^16^ (**Figure 3b**).

**Figure 3.**
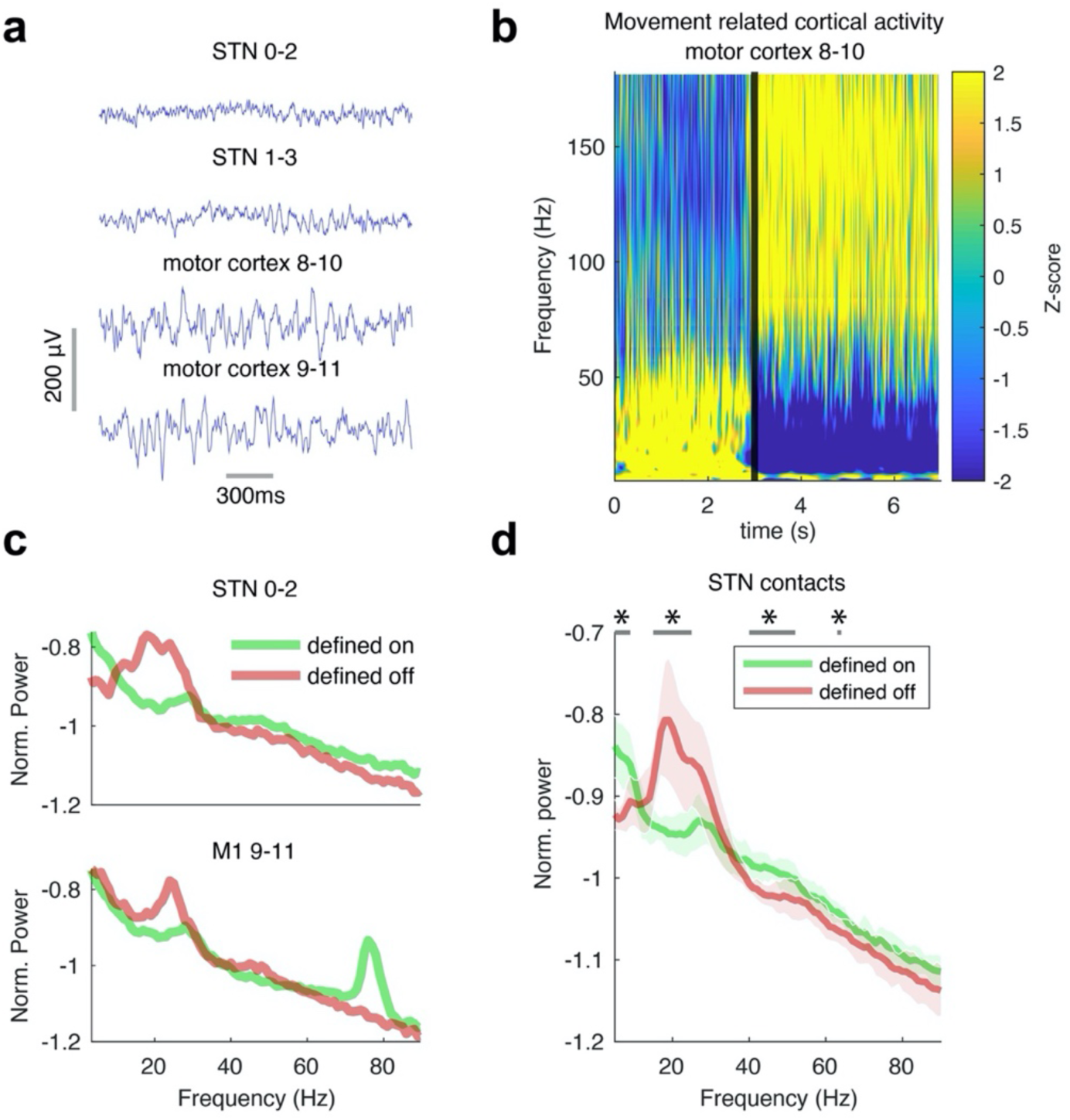
Data examples and demonstration of effects of movement and levodopa in-clinic. **a**, Example field potentials recorded from right hemisphere, cortical (top) and STN (bottom). Horizontal grey (bottom left) line represents 300ms, vertical line is 100 uV. **b**, Example spectrogram of ECoG activity (bipolar recordings contacts 8-10) showing canonical movement-related alpha-beta band (8-35 Hz) decrease, and broadband (50-200 Hz) increase, consistent with placement over sensorimotor cortex (from RCS04), recorded 27 days post-implantation (sampling rate 500 Hz). Dotted vertical line is the onset of movement. Color scale is z-scored. **c**, Example power spectra of STN and motor cortex field potentials showing oscillatory profile of off-levodopa (red) and on-levodopa (green) states (patient RCS01), from 30 second recordings. **d**, STN beta amplitude is consistently reduced in the on-medication state. (p<0.001). Average PSD plots across both hemispheres, both recording montages, and all 4 patients (average – dark line, shaded area is one standard deviation). Horizontal bar shows frequency bands that had significant differences between states. Antiparkinsonian dopaminergic medications such as levodopa are thought to induce profound changes in oscillatory activity in the basal ganglia, based on previous perioperative recordings in humans using externalized brain leads^2^. Thus, we collected data in defined medication states: after 12 hour withdrawal of antiparkinsonian medications (“off”), and after a suprathreshold dose of levodopa/carbidopa (“on”). The on-medication state was associated with the expected reduction in subthalamic beta band (13-30 Hz) activity in each individual case (**Figure 3c**) and across all 16 subthalamic recordings (p < 0.001, Generalized Estimating Equations (GEE)) (**Fig 3d**). Thus, acute effects of levodopa observed previously with externalized leads were validated here by wireless data transmission from an implanted device, recorded 2-4 weeks after surgery. In patients with prominent dyskinesias, a motor cortex oscillation at 60-90 Hz appeared in the on-state^12^ (**Figure 3c**).

### Physiological signatures of motor signs identified by pairing at-home neural recordings with wearable monitors

A challenge in the field of invasive human brain recordings has been validation of findings from acute in-hospital recording paradigms, in chronic settings as patients go about activities of daily living. Here, patients streamed eight channels of neural data at home over a total of 548 hours, while experiencing their typical on/off fluctuations from their habitual schedule of antiparkinsonian medications, as well as during sleep. Data were collected 2-4 weeks after device implantation, prior to initiating standard clinical neurostimulation. Clinically validated wrist-mounted wearable monitors worn bilaterally (Parkinson’s KinetiGraph (PKG) watch)^17^ provided numerical scores for bradykinesia and dyskinesia every 2 minutes based on a 10-minute moving average (**Figure 4a**). All patients experienced motor fluctuations as evident from their PKG watch data with periodic variations in scores similar to **Figure 4a**.

**Figure 4.**
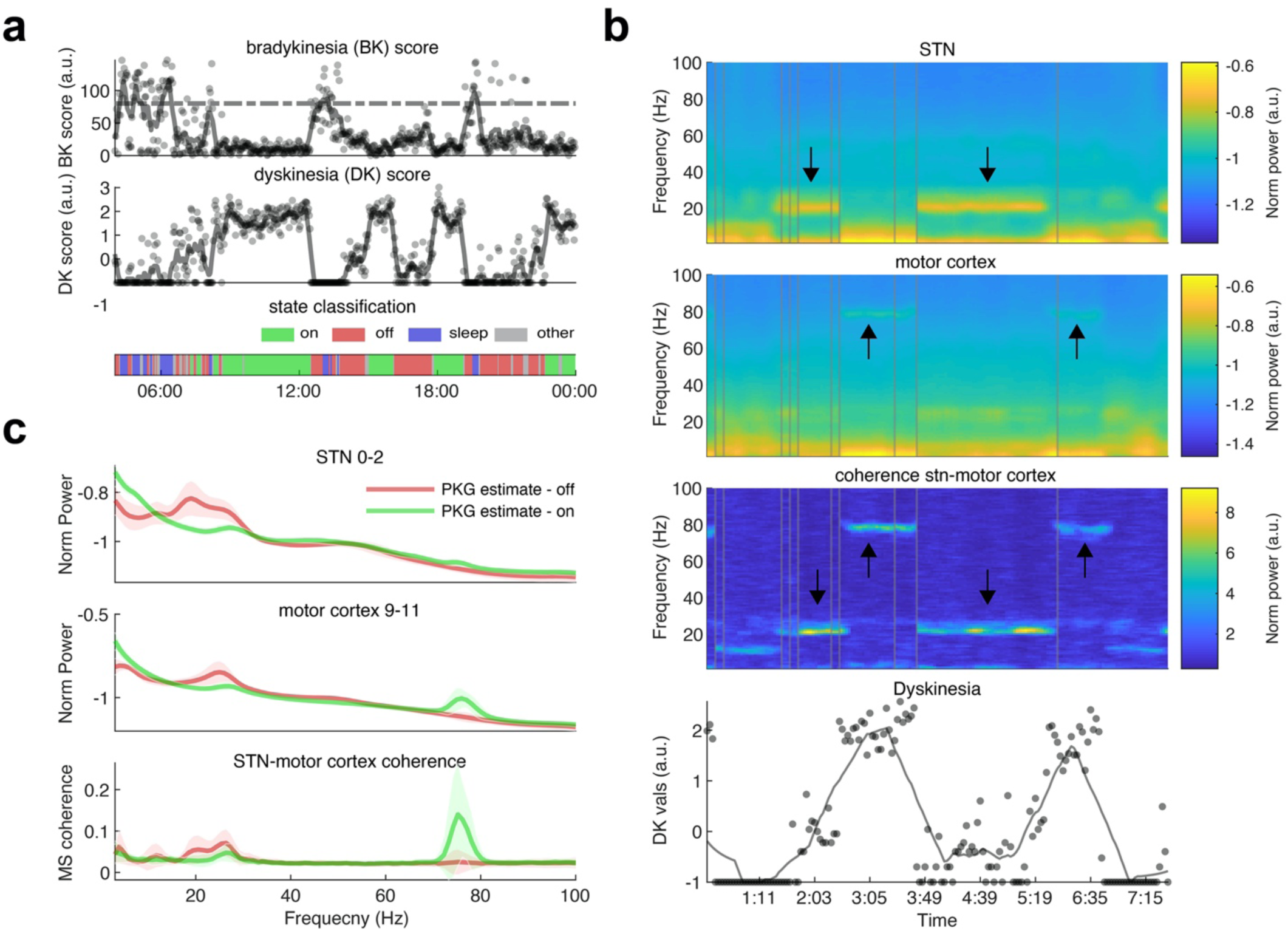
Decoding motor fluctuations from long duration recordings at home, single subject example (RCS01). **a**, Data from wearable Personal KinetiGraph (PKG) monitor reports scores for bradykinesia and dyskinesia in 10 minute intervals. Example from 1 day. Assignment of state is shown in the lower bar. **b**, Capturing transitions between immobile (off) and mobile/dyskinetic (on) states. Top, spectrograms for STN and motor cortex, and STN-motor cortex coherence over a 7.5 hour period (all times PM). Arrows indicate frequency bands sensitive to on-off fluctuations. Grey vertical lines show areas where the recording was discontinuous and was concatenated. Bottom, PKG dyskinesia scores indicate four transitions between off (low dyskinesia) and on (high dyskinesia) states. These are associated with transitions in beta and gamma oscillatory activity. **c**, Power spectra of STN and motor cortex, and STN-motor cortex coherence for all awake data from patient RCS01, segregated by mobile and immobile states (categorized by PKG) and averaged. Shaded error bars represent one standard deviation

Neural data were analyzed in 10 minute segments, to correspond to the length of time over which motor scores were calculated from the PKG wearable device. A power spectrum was calculated for each 10 min segment and superimposed for all single site recordings within subjects (**Figure S1**). Sleep states, as identified by PKG scores, typically showed prominent delta activity in both STN and cortex (**Figure S1**). For awake data, the power spectra calculated from 10 minutes segments were then segregated into mobile (“estimated on”) and immobile (estimated “off”) segments by PKG scores and averaged. A time-frequency analysis over a single day (7.5 hours) from a single subject shows that transitions between on and off states are associated with simultaneous rapid transitions in beta and gamma band oscillatory activity, as well as in in coherence between STN and cortex (**Figure 4b)**. Averaged LFP power spectra segregated by the wearable monitor (42.7 hours of recording) broadly recapitulates the patterns discerned in defined on/off states in-clinic: A prominent STN beta band peak when off, which disappears in on states, and a prominent motor cortex gamma band peak at 75 Hz when on with dyskinesia (**Figure 4c**, compare to brief in-clinic recordings in **Figure 3c**).

STN-motor cortex coherence also distinguished on and off states (**Figure 4c**), suggesting the use of basal ganglia-cortical oscillatory interactions, in addition to single-site oscillatory profiles, for the identification of motor function from neural recordings. Variations on these spectral patterns in other subjects occurred, consistent with cross-subjects variations in their most prominent motor signs (**Figure S1**). For example, in the one subject with prominent off-period tremor (RCS03), an oscillation at twice tremor frequency appeared, especially in STN-cortical coherence, during tremor episodes (**Figure S1)**. This physiological signature of tremor has been detected previously in the cortex using magnetoencephalography^18^.

The number of hours of data streamed at home by each patient is in **Figure 5a**. Across all subjects, the sites and frequencies of significant separation in neural data between estimated on and estimated off states, occurred in the STN beta band and cortical gamma band (**Figure 5b**).

**Figure 5.**
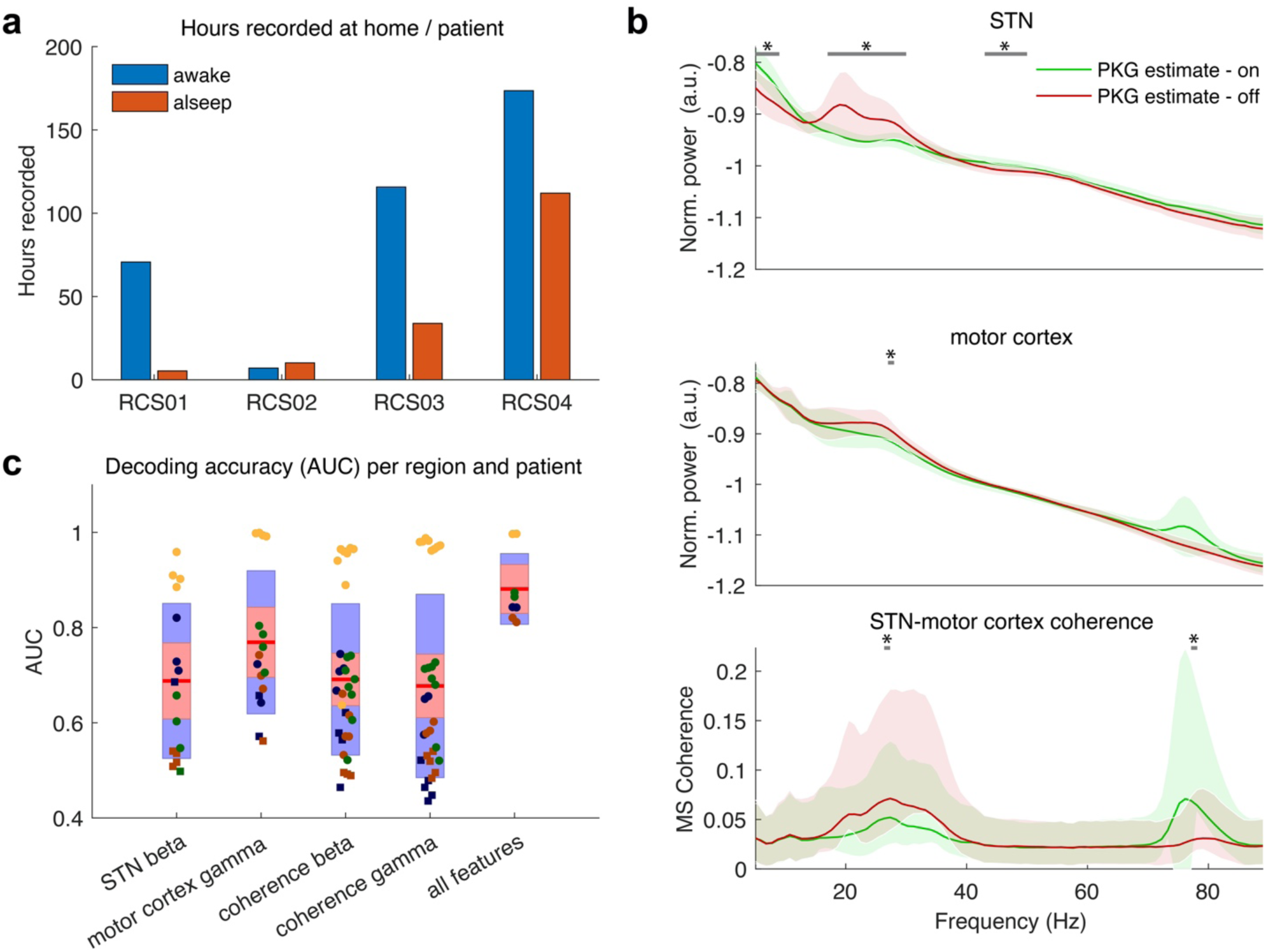
Decoding motor fluctuations from long duration recordings at home, group data. **a**, number of hours recorded by each patient **b**, Power spectra of STN and motor cortex, and STN-motor cortex coherence, for home recordings combined across all subjects. Horizontal bars with asterisks indicate frequencies that differed between states after correction for multiple comparisons. **c**, Area under the curve (AUC) from ROC (receiver operator curve) analysis, showing that utilizing data from both STN and cortex is better able to discriminate mobile and immobile states (as segregated by PKG scores), than either site alone. The input for the computation of each ROC curve (a single point on graph) was calculated by computing average beta band spectral power from STN (within each patient, side and two possible contact pairs – yielding 16 points), average gamma power from cortex, or a combination of estimates across recording montages (two values from STN and two from cortex). The four colors correspond to the four study subjects.

The specificity and sensitivity of these neural features for decoding on and off behavioral states was quantified across all recordings in all subjects using receiver operating characteristic curve (ROC) analysis (**Figure 5c**). Discriminating on and off states was possible from either STN beta activity (Area Under Curve (AUC) range 0.5-0.96), or motor cortex gamma activity (AUC range 0.56 -1). An important debate in the area of decoding behavior state from brain recordings, is whether to utilize subcortical data, cortical data, or both. The combination of subcortical and cortical data provided the best behavioral discrimination (AUC range 0.81-1.0) (**Figure 5c**), underscoring the utility of *multisite* sensing for bidirectional interfaces. The significance of decoding was tested within subjects non-parametrically by shuffling the “on” and “off” labels and repeating the 5-fold cross validation. Single site spectral power beta or gamma bands, in ten minute recordings, was predictive of motor state in most but not all hemispheres, whereas when both recordings sites were combined and cortex-STN coherence was included, ten minute segments of neural data significantly segregated on and off states for all hemispheres (**Figure 5c**).

### Unsupervised clustering of neural data for identification of distinct behavioral states

A major goal for clinical utilization of bidirectional interfaces in neurology and psychiatry is the implementation of adaptive DBS, using electrophysiological signatures (biomarkers) of specific signs and symptoms^4,15,19^. To accomplish this efficiently when biomarkers are highly individualized, distinct brain states should be identifiable using “unsupervised” clustering algorithms, with which neural data are categorized without the need for paired behavioral monitors, patient reporting, or clinician ratings. We employed an unsupervised clustering method based on density of data points in dimensionality-reduced power spectra^20^ (“density-based clustering”, **Figure 6a**). We chose this method since it does not require defining the number of states (unlike k-means clustering). Indeed, unsupervised clustering algorithms were able to identify clusters that corresponded to supervised clustering (on/off state estimates clustered by wearable PKG monitor) with high concordance (mean 74%, range 59%-89% for density based clustering, mean 74.5% range 59%-90% for template based clustering)(**Figure 6c**). A second clustering method leverages the power spectra of brief in-clinic recordings, done in defined behavioral states, as “templates” for unsupervised clustering in home data (“template based clustering”, **Figure 6b**). In disorders for which behavioral states are readily induced in-clinic, this is a powerful method to identify those states in home recordings. Template clustering also shows high concordance with supervised clustering for STN data (**Figure 6c**).

**Figure 6.**
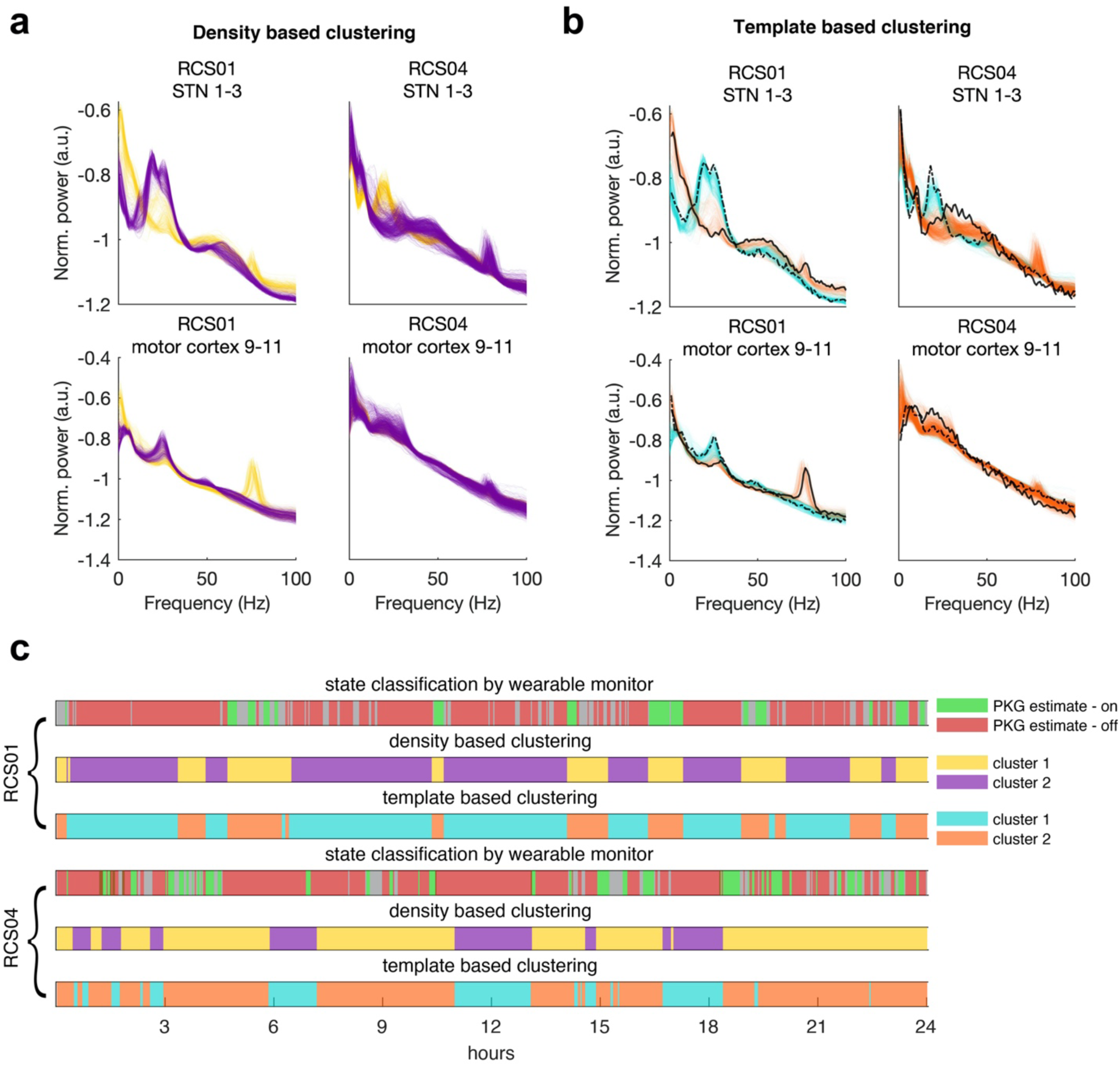
Unsupervised clustering segregates neural data into specific behavioral states. Examples patients are RCS01 and RCS04. All raw data (recorded in the awake state) were segregated using unsupervised clustering algorithms with two different paradigms: **a**, Unsupervised clustering using the density based method of Laio^20^. **b**, Clustering of PSDs based on template PSDs from in clinic recording in defined on/off medication states. Black lines are the template PSD’s (dotted = off medication, solid = on medication). **c**, Concordance between brain states derived from unsupervised and supervised clustering methods (24 hour data sample, STN only). Barcodes compare motor state estimates derived from the wearable monitors, with the clusters derived from type of clustering algorithm.

### Advanced device features that enable embedded adaptive stimulation

A central challenge in the design of implantable bidirectional neural interfaces is the enabling of sensing from the same array that is used for therapeutic stimulation, a critical capability for adaptive neurostimulation. This is particularly challenging with subcortical targets, since the amplitude of field potentials from non-layered structures is often <10 µV, which is 5- to 10-fold less than those recorded from the cortical surface. Previous bidirectional interfaces have usually allowed meaningful sensing only during periods when stimulation was absent, or required sensing from a distant electrode array, such as from a cortical array during subcortical stimulation^21^. Here, using a “sandwiched” sensing configuration to take advantage of common mode rejection (bipolar sensing from the contacts on either side of a monopolar stimulating contact), we show that STN beta power is reduced by chronic therapeutic stimulation, without a concomitant change in cortical beta power (**Fig 7a**). Violin plots show how chronic stimulation affects the variability of STN beta band activity (**Fig 7b**). Prior to therapeutic stimulation, the range of beta amplitudes shows a bimodal distribution, corresponding to on- and off-states. Chronic therapeutic DBS eliminates pathologically elevated beta epochs, but preserves variability of beta band activity within a smaller range. Preservation of task-related variation in beta activity may be important for normal movement during DBS^22^.

**Figure 7.**
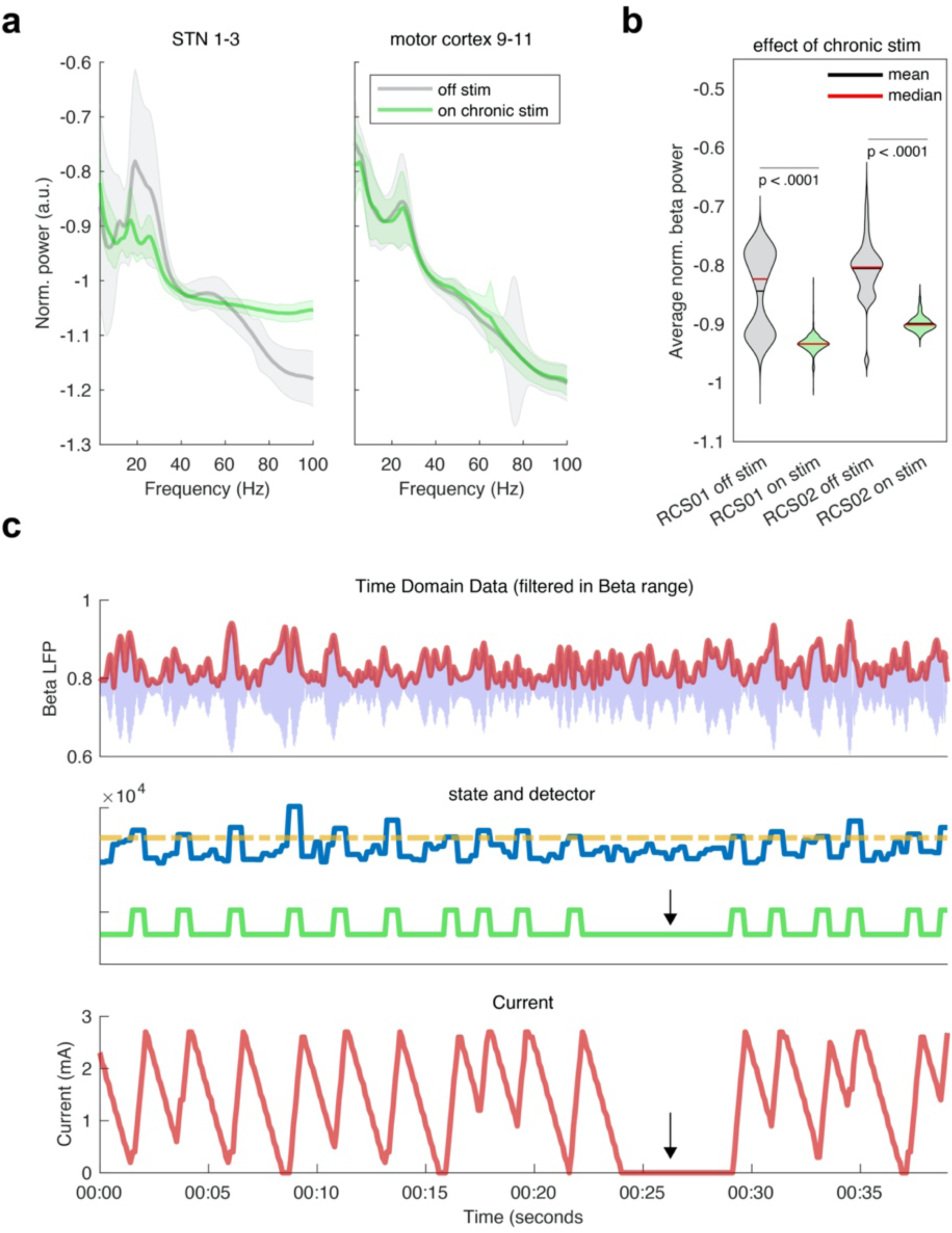
Recording during therapeutic DBS, and demonstration of adaptive DBS. **a**, Chronic recording from same quadripolar STN contact array as utilized for therapeutic stimulation. Example from RCS01. Average PSDs for 10 min data segments segregated by off stimulation (grey), and on stimulation (green), over a total of 87.5 hours of recording at home. **b**, Violin plots showing the average beta power (5Hz window surrounding peak) off/on chronic stimulation in two rigid/akinetic patients. Chronic STN DBS eliminates periods of pathologically elevated STN beta band activity, but preserves variability within a smaller range (median off stimulation versus chronic stimulation differ**). c**, Example of embedded adaptive DBS algorithm designed to trigger off of beta bursts. Top panel is filtered STN time domain data (blue) in the beta range (17-19Hz) centered around subject specific beta peak with amplitude (Hilbert transform, red) superimposed. Middle panel – threshold chosen for algorithm in (dotted orange line) and the detector (blue) which is a smoothed estimate of instantaneous beta power. Algorithm state is in green such that when the detector crosses the line state is “high” and when it is below the line state is “low”. Bottom panel – current amplitude in mA. This algorithm is designed to “trim” beta bursts as advocated by Little et. al. (2013). When a beta burst is detected stimulation is rapidly ramped up followed by rapid decrease when burst subsides. Arrow indicates a 5 second epoch during which the detector is not triggered.

A promising but technically demanding approach to adaptive DBS is to use very short trains of DBS triggered on bursts of oscillatory activity, to shorten the duration of pathologically prolonged bursts^23^. This approach has shown efficacy in PD that is greater than that of standard continuous stimulation^10^, but has only been implemented using externalized leads connected to external amplifiers and computers, or with implanted devices streaming to an external computer in a “distributed mode”^24^. We therefore tested this strategy for adaptive DBS, in a fully embedded paradigm (**Figure 7c**). A common pitfall of feedback control systems is the occurrence of “limit cycles”, that is highly regular oscillatory behavior in which each state change triggers the opposite change, such that the detector fails to respond appropriately to neural activity changes. Irregular changes in detector state show appropriate algorithm performance without converging to a limit cycle (arrow in **Figure 7c**).

## Discussion

Here we report the first human use of an implanted bidirectional neural interface designed for continuous wireless streaming of neural data for long periods, at home during normal daily activities. In four PD patients implanted bilaterally with both motor cortex and basal ganglia leads, we show that patterns of oscillatory activity in both structures can decode hypokinetic and hyperkinetic states, as demonstrated by pairing the recordings with wearable monitors which track these motor fluctuations behaviorally. We show that brain states are also separable by an unsupervised clustering algorithm applied to the power spectra of short data segments, providing a rapid method for personalized biomarker discovery. Finally we demonstrate technical device capabilities that have not previously been readily achievable in implantable sensing devices: sensing during stimulation from adjacent contacts of a multipolar array, and embedded adaptive DBS utilizing small subcortical signals for control.

### Circuit discovery in humans using field potentials

Analyses of field potentials from invasive recordings in humans have shown great promise for discovering the circuit basis of a variety of brain disorders^25^. Field potentials represent the summed, synchronized activity of a neuronal population near the recording contact and usually have a strong oscillatory component, thus offering an excellent probe of neural synchronization. In normal function, oscillatory synchronization provides a mechanism for flexible communication between functionally related brain regions, as elucidated in the “communication through coherence” hypothesis^26^. Many brain disorders are now thought to be related in part to abnormalities of oscillatory synchronization^25^. In patients undergoing invasive monitoring for epilepsy, mood and anxiety have been decoded from patterns of oscillatory synchrony in field potential recordings^27,28^, and this approach shows promise in major depression^29^ and in obsessive compulsive disorder^30^. These findings have generated great interest in utilizing long term brain recordings from implantable devices in a wide variety of disorders^3^. Identification of neural signatures of disease states translates to improved therapies in several ways: 1) detecting circuit abnormalities that predict response to neurostimulation therapy, 2) optimizing symptom-specific stimulation algorithms, and 3) the use of these neural signatures for adaptive neurostimulation.

Here, we utilized chronic home recordings to inform the “oscillation hypothesis” of movement disorders pathophysiology. A seminal observation, made using perioperative recordings from externalized leads, was that the amplitude of beta band oscillatory activity in the basal ganglia is relatively elevated in parkinsonian states^31,32^, and its attenuation is a reliable index of the effects of therapeutic intervention (both oral levodopa^33^ and DBS^31,34^). As a result, the “oscillatory synchrony” hypothesis of parkinsonian akinesia^2^ and bradykinesia has begun to replace the earlier “rate model” of parkinsonian pathophysiology^35^. However, this hypothesis has only been tested in non-naturalistic clinical situations in severe and controlled off- and on-medication states, which might not reflect neural activity in the real world during normal activities of daily living. Here, we show that subthalamic beta band activity can indeed consistently track the severity of motor fluctuations in PD in chronic, real-world use. However, motor cortical gamma band (65-90 Hz) oscillations, and basal ganglia-cortex coherence in beta and gamma bands, also track motor signs. In many cortical regions, gamma oscillations coordinate local neuronal ensembles during task performance^25^. Our findings suggest that an imbalance in basal ganglia beta band activity and cortical gamma activity may be critical for the expression of motor signs of PD, and highlight the utility of multisite recordings within a network, both for optimal circuit discovery and for high fidelity characterization of brain states.

The analysis of large amounts of neural data “supervised” by wearable monitors is labor-intensive and depends on the accuracy of the wearable monitor as well as a prospective understanding of the behavioral states that need to be tracked. Unsupervised clustering, in contrast, provides efficient identification of individualized neural signatures of disease state from large data sets, does not depend on patient compliance with a wearable monitor, and may identify previously unrecognized patterns of neural activity to generate new hypotheses about circuit mechanisms and biomarkers. Here, unsupervised clustering was able to distinguish brain states corresponding to on- and off-periods, similar to those identified with wearable monitors. (**Figure 6**). These methods may prove especially important for identifying neural correlates of psychiatric states, for which wearable monitors for objective tracking are less advanced than in movement disorders. Unsupervised clustering can also be supplemented by the use of neural activity “templates” derived from brief in-clinic recordings in a defined behavioral state, to objectively search for the occurrence of those states during chronic neural recording at home (**Figure 6b**). Whereas the templates in our study were provided by defined medication on and off states, in the application of this method to psychiatric illness, templates could be established by “provocative” tests such as brief exposure of a patient with obsessive-compulsive disorder to images that trigger obsessions^36^.

### Second generation bidirectional interfaces: new capabilities and their implications

Several fully implantable brain devices that combine neural sensing with therapeutic neurostimulation have been introduced prior to the present work. The RNS device (Neuropace) is clinically approved in Europe and USA for contingent stimulation in some types of epilepsy^4^. It has been used for sensing and adaptive DBS in Tourette’s syndrome in an investigational study^37^. A first generation precursor of RC+S, Activa PC+S (Medtronic), has been used under investigational protocols for brain sensing in PD^6,12,38-40^, essential tremor^41^, epilepsy^42^, pain^43^, and locked-in syndrome^44^. However, both devices are designed for brief recording rather than continuous streaming of time series data, are problematic for recording during therapeutic stimulation on adjacent contacts of an array^38^, and are inflexible with respect to the type of adaptive DBS algorithms that can be embedded^3^. While earlier devices have allowed long term streaming of spectral power in a predefined frequency band^44^, time series data are important since they do not require a priori knowledge of the most relevant frequency bands, and allow the exploration of many other metrics of synchronization including phase coherence^12^, phase-amplitude coupling^45^, and waveform shape^46^.

Second generation bidirectional interfaces require several technical and conceptual innovations to overcome the limitations of first generation devices. One important innovation is allowing academic investigators full access to a device’s Application Programming Interface^7^. Thus, for the first time, researchers may write their own code to control the research functions of the device, including home data streaming and adaptive DBS. This provides great flexibility to tailor device functions to specific studies. Here, we wrote a patient-facing graphical user interface customized for PD patients, for data streaming at home from bilateral devices and for marking relevant data such as timing of medications. This is readily usable by patients without the presence of study investigators, and provides relatively seamless re-initiation of streaming whenever the patient was in range of the receiver (12 m). Thus, patients were able to use this for many hours of continuous data streaming, while freely moving and untethered to receivers. The rechargeable battery obviates potential concern about the need for surgical replacement of the pulse generator in the setting of energy-intensive data streaming at home. Artifact management strategies, including active recharge for biphasic pulses and customizable signal blanking during therapeutic stimulation, allow sensing of neural biomarkers during therapeutic stimulation to a much greater extent than prior devices^38^.

The capabilities of second generation neural interfaces open up new areas for basic science investigations and for clinical translation. Neurophysiological investigations in humans have classically been done during brief repetitions of well defined, but artificial, tasks, in laboratory or clinical environments. The capacity for high volume data streaming for long periods of time in the real world allows, for the first time, study of human neurophysiology in naturalistic environments^47^. The statistical power offered by high volume longitudinal data is based on within-subjects comparisons of neural activity over many exacerbations and remissions of specific signs and symptoms of disease states, supplemented by wearable monitors to independently categorize behavioral state. Spontaneous fluctuations in normal behaviors could be studied using the same approach.

#### Potential for Adaptive DBS

While continuous neurostimulation is now employed in many conditions, it may induce adverse effects such as hypophonia^11^ or dyskinesia^12^ in PD, mania in OCD^48^, or seizures from cingulate stimulation for pain^43^. Efficacy of continuous stimulation may also suffer from waning effectiveness, such as in chronic pain^49^. If stimulation were only delivered contingent on the relevant patterns of abnormal circuit activity, it could respond to changing brain needs and reduce adverse effects^15,19^. Further, adaptive DBS has the potential to shape neural circuits, such as shortening of pathologically lengthened subthalamic beta bursts in PD^22^, so as to be more effective than open loop, continuous DBS. The present study demonstrates this capability in an embedded paradigm within a fully implanted device, whereas prior demonstrations of adaptive DBS in PD triggered on small subcortical beta bursts, have only been possible using externalized brain leads^10^ or using first generation implanted devices in “distributed mode” (streaming to an external computer) in hospital settings^24^.

### Challenges and future developments

A new challenge of RC+S, inseparable from its high flexibility, is the requirement for academic investigators to write their own software to control its functions, and to document its compliance with FDA requirements. Investigators thus need to hire or contract with a dedicated software engineer familiar with medical devices, or collaborate closely with other groups that do so. Here, this complexity was mitigated by establishing a multi-institutional collaborative environment in which device control software and regulatory templates are freely shared between academic groups (https://openmind-consortium.github.io). In the future, use of neural interfaces during normal activities could be facilitated by allowing streaming directly from the implanted device to smartphones. Smartphones may also provide applications for behavioral monitoring that could then be readily paired with neural recordings^50^.

## Methods

### Inclusion criteria and clinical characterization

Four study subjects were recruited from a population referred for implantation of deep brain stimulators for PD. Subjects were evaluated by a movement disorders neurologist and met diagnostic criteria for PD^1^. Baseline motor function was evaluated using the Movement Disorders Society (MDS) Unified Parkinson’s Disease Rating Scale (UPDRS), parts I-IV. The motor subscale (UPDRS-III), was rated both “off” (12 hour after withdrawal of antiparkinsonian medication) and “on” (after a supratherapeutic dose of levodopa/carbidopa). Patients were evaluated by a neuropsychologist to exclude significant cognitive impairment or untreated mood disorder. Inclusion criteria were: Motor fluctuations with prominent rigidity and bradykinesia in the off medication state, baseline off-medication UPDRS-III scores between 20 and 80, greater than 30% improvement in UPDRS-III on medication compared to off medication, and absence of significant cognitive impairment (score of 20 or above on Montreal Cognitive assessment). The study was approved by the hospital institutional review board (IRB) under a physician sponsored investigational device exemption (IDE), protocol # G180097. The study was registered at Clinical Trials.gov (NCT03582891). Patients provided written consent in accordance with the IRB and the Declaration of Helsinki. The full IDE application and study protocol have been shared with other researchers via the Open Mind initiative (https://openmind-consortium.github.io).

### Surgery, device models, and lead localization

All patients underwent bilateral placement of cylindrical quadripolar deep brain stimulator leads into the subthalamic nucleus (Medtronic model 3389, 1.5 mm contact length and 2.0 mm intercontact spacing), bilateral placement of paddle-type quadripolar cortical paddles into the subdural space to cover precentral gyrus (Medtronic model 0913025, 4 mm contact diameter and 10 mm intercontact spacing), and bilateral placement of investigational sensing implantable pulse generators (IPGs) in a pocket over the pectoralis muscle (Medtronic Summit RC+S model B35300R). The IPG and leads were connected by 60 cm lead extenders (Medtronic model 37087), two on each side (**Figure 1**). STN leads were initialized as contacts 0 to 3 (0 is the deepest contact)(**Figure 2a**). Cortical leads were initialized as contacts 8 to 11 (8 is the most posterior contact)(**Figure 2c**).

The surgical technique for placement of permanent subdural paddle leads during DBS implantation surgery was previously described in detail^2^. Briefly, the paddle lead was placed in the subdural space through the same frontal burr hole used for the subthalamic lead. At least one contact covered the posterior precentral gyrus (presumed primary motor cortex), approximately 3 cm from the midline on the medial aspect of the hand knob. Adequate localization of the ECoG strip was confirmed using intraoperative CT computationally merged to the patient’s preoperative MRI^3^ (Stealth8 Cranial software, Medtronic Inc.) Functional localization of the ECoG strip was verified by reversal of the N20 somatosensory-evoked potential from median nerve stimulation (**Figure 2b**). The exiting wire from the cortical contact array was secured to the skull with a titanium miniplate.

The subthalamic lead was placed using frame-based stereotaxy and confirmed by microelectrode recording in the awake state using standard methods^4^. Proper location in the motor territory of the STN was verified by eliciting movement-related single-cell discharge patterns. The DBS lead was placed with the middle two contacts in the dorsal (motor) STN, the most superior contact dorsal to the STN, and the most inferior contact in ventral STN (**Figure 2a**). The free ends of the cortical and subthalamic leads were coiled under the ipsilateral parietal scalp. The remaining hardware was placed under general anesthesia. The free ends of the cortical and subthalamic leads were connected to the lead extenders, which were tunneled down the neck to the IPG. Each IPG was connected to the ipsilateral cortical and STN leads. Medical adhesive was placed at the junction of the lead extenders and IPG to reduce contamination of neural signals by EKG artifacts. Two months postoperatively, locations of leads were again verified, by postoperative CT computationally merged to the patient’s preoperative MRI using Stealth8 Cranial software (**Figure 2c**).

### RC+S device characteristics and programming

The Summit RC+S is an investigational rechargeable bidirectional neural interface that offers the researcher a great degree of flexibility through access to the device’s application programming interface (API)^5,6^. It is a 16-channel device that can simultaneously stream four bipolar time domain channels (250/500Hz) or two channels at 1000Hz. It can simultaneously provide standard therapeutic stimulation on up to two quadripolar leads, and can also perform adaptive deep brain stimulation using algorithms programmed on the device (“embedded” mode) or algorithms on an external computer, through wireless communication (“distributed” mode). In addition to voltage time series data, RC+S can stream up to 8 predefined “power channels” (spectral power within a predefined frequency band), event markers, stimulation parameters, embedded adaptive DBS performance parameters, and triaxial accelerometry from an embedded accelerometer.

For all research functions including configuring and initiating sensing, and embedded or distributed adaptive DBS, investigators control the device by writing software in C# within the device API, accessed using a “research development kit” (RDK, Medtronic model 4NR013) provided by the manufacturer. We wrote two graphical user interface (GUI)-based applications to configure and initiate streaming data from one or two RC+S devices simultaneously. One application is used by the research team and allows configuration of sensing parameters and streaming data in-clinic. The other application is “patient facing” and contains a simplified application that allows the patient to control streaming in a home environment and report symptoms or medications taken, Applications rely on a dynamic linked library (DLL), supplied by Medtronic, Inc., that is specific for Microsoft Windows operating systems and Intel processor platform. The DLL provides the API to investigators and is not compatible with streaming data to mobile devices. Both applications are available at https://openmind-consortium.github.io. We wrote and documented software in compliance with FDA code of federal regulation CFR 820.30, which specifies design controls for implantable human devices. **Figure 1** provides a schematic of the data streaming configuration.

### In-clinic data recording in defined on/off states

STN and cortical field potentials were sampled at 500 Hz in clinic three weeks postoperatively in both “on” and “off states (during which levodopa medication was withdrawn for at least 12 hours), and a movement disorders neurologist administered the UPDRS-III rating scale in both states. The three week time point was chosen to allow recovery from the “microlesion” effect of lead insertion^7,8^, but prior to initiating chronic therapeutic stimulation at 1 month after surgery. Recordings were done at rest, and during a binary choice iPad reaching task which has been previously described^9^. Rest recordings were one to two minutes long, and iPad reaching task recordings were 3-5 minutes long.

### At home data streaming paired with wearable monitors

Patients initiated home recordings using the patient-facing GUI on a Microsoft Surface Go computer with broadband cellular service (weight 1.15 pounds, dimensions – 245.00 × 175.00 × 8.30mm (height, width thickness)). We provided this computer to each study subject along with training in its use. Streamed data contained no personal health information (PHI). Data were encrypted and uploaded to a secure cloud environment operated by UCSF. Patients were able to use the application to report medications taken and to rate their motor signs. The application automatically connects to the device when patients are in range (approximately 12 meters). Patients collected data in 1-2 week recording “sprints” in which they were instructed to carry the computer with them and stream continuously if possible.

The summit RC+S system uses a user datagram protocol (UDP). Data are transmitted in discrete intervals or “packets” of variable duration averaging 50 ms. Occasional data packets are lost and these “dropped packets” must either be interpolated or discarded. This can also be partially mitigated by changing sampling rates and using a lower data streaming rate bitrate. Failure of packet transmission (“dropped packets”) occurred for 1-5% of packets, even when patients were in range of receiving devices. We have written specialized software in Matlab to account for dropped packets which is available on our GitHub page https://github.com/starrlab/rcsviz.

Data are time stamped using the pulse generator clock time. Data were recorded at 250Hz in at the patient’s home, lower than the in-clinic rate of 500 Hz. Four time domain channels were streamed using a bipolar recording configuration in which we verified adequate signal during a montage recording obtained one to two days postoperatively. Patients also streamed actigraphy at 64Hz from the embedded accelerometer, event related information and power channels. The Summit RC+S device has several configurable device filters that must be chosen. All filters are applied after digitization, and low pass filters are applied twice – before and after amplification. In the absence of therapeutic stimulation, we used a high pass filter of 0.85Hz and low pass filter of 450 Hz before amplification and 1700Hz after amplification.

To track parkinsonian motor signs at home, for subsequent correlation with paired neural data, patients wore wristwatch style monitors (Parkinsons’s KinetiGraph System (PKG), Global Kinetics Inc)^10,11^. The PKG reports scores for bradykinesia, dyskinesia and tremor every two minutes as well as the timing in which medication were taken. The PKG uses a 3 axis accelerometer (similar to that embedded in RC+S) to assess parkinsonian symptoms according to a proprietary commercial algorithm that has been validated. We confirmed accurate temporal synchronization between the PKG and the RC+S by comparing their actigraphy data.

### Maintaining continuity of data streaming

Benchtop tests show that streaming of four time domain channels at 500Hz while providing therapeutic stimulation, can be done for approximately 30 hours before recharging the RC+S pulse generator. However, several factors may reduce the duration of continuous streaming. The CTM relay device (**Figure 1**), as supplied by the manufacturer, is powered by two AAA batteries that only last for 4-5 hours during streaming. We therefore developed an external “battery pack” for the CTM that extends this range to 12 hours. Both RC+S pulse generators can be recharged simultaneously in 30 minutes. For most home streaming we used lower sampling rates (250Hz) than the device allows, in order to cap the bitrate at 4500 bits/second but allows for longer range and fewer dropped packets. Our patients wore a specialized vest with pockets for the CTM so that it remained in close proximity to the DBS device to avoid frequent packet drops.

### Therapeutic continuous stimulation and recording during stimulation

To implement standard DBS therapy, clinicians are provided with a tablet programmer (Medtronic model 4NR010) to allow setting parameters for standard continuous therapeutic stimulation, and a patient programmer (Medtronic model 4NR009) that allows the patient limited control over some parameters under limits set by the clinician. One month after implantation, study clinicians began programming the STN lead(s) to achieve the best clinical result. The cortical lead was never used for stimulation. Clinicians attempted programming using monopolar mode from one of the middle two contacts (contacts 1 or 2), as these montages are compatible with bipolar sensing in a “sandwiched configuration” around the stimulating contact. Sensing from STN leads programmed to stimulate in a bipolar mode or programmed with a stimulation montage that includes contacts 0 or 3, precludes STN sensing during stimulation because of excessive stimulation artifact. For some subjects, 80 hours of data were streamed at home during therapeutic stimulation, to document effects of stimulation on STN and cortical field potentials. Both on-device low pass filters were set at 100Hz for STN recordings during stimulation to limit stimulation artifacts). Sense blanking was set at 0.33ms

### Data extraction and management of lost data

Data streamed to the researcher or patient facing applications are assembled into JavaScript Object Notation (.JSON) format. This is a light-weight data-interchange text-based format that in RC+S merges meta-data and actual data across several different file types distributed. Data are polled from the device in configurable intervals (50 ms in our case) in first-in, first-out (FIFO) fashion. The IPG has a 16 bit clock-driven tick counter that rolls over every 6.553 seconds (least significant bit (LSB 100 microseconds)). This can be combined with an estimate of system time (LSB seconds) to accurately account for lost packets. We wrote Custom software in Matlab to extract the data from the .JSON format and discard packets that have corrupted data. The extraction code is available on GitHub (https://github.com/starrlab/rcsviz).

### Data Processing (in-clinic data)

Data were inspected for artifacts and contiguous 30 second data segments for which no packet loss occurred were chosen for analysis. We calculated power spectral density (PSD) using the Welch Method in Matlab (pwelch, 500ms window, 250ms overlap) from 4 contact pairs (0-2, 1-3 in STN and 8-10, 9-11 in the cortical lead) per implant side. This yielded 8 PSD’s per patient in the on and off medication condition. For each PSD we computed the average beta signal (13-30Hz) subcortically or gamma (65-85Hz) cortically across patients. We compared the average frequency specific spectral power on/off medication response using generalized estimating equations (GEE) to account for non-independence of recording locations within patients^12^ using the GEEQ toolbox for Matlab. For movement-related changes in spectral power, data were filtered using a two way FIR1 filter (eegfilt from eeglab toolbox with fir1 parameters) in frequencies between 1-200Hz. Data from all trials were aligned relative to the onset of movement (time 3) and averaged. The averaged amplitude was normalized by a 1000ms window prior to cue presentation (time 0). Data were z-scores by subtracting the average baseline amplitude and dividing by the baseline standard deviation. This z-score procedure was performed for each frequency separately.

### Data Processing (home data)

Data were divided into 30 second contiguous chunks in which no packet loss occurred. We calculated power spectral density (PSD) using the Welch Method in Matlab (pwelch, 250ms window, 125ms overlap). Data were averaged in the power domain between 40-60Hz (range selected to avoid frequency bands of physiological interest) and outliers larger than 2 standard deviations were excluded from further analysis. This was usually due to the presence of transient artifacts in the data and only affected 1.3% on average (range 0.1-3.1%) of the data. Data were normalized by dividing each PSD by the average power between 3 and 90 Hz. In addition to the PSD, the magnitude squared coherence was also computed for the four possible contact pairs each recording montage (mscohere in Matlab, 256 ms window, 50% overlap, 256 discrete Fourier transform points).

### Synchronization of neural data with wearable monitor state estimates

The RMS voltage of the internal built-in accelerometer was used in order to verify the synchronization between the PKG and the RC+S data. The (root mean square) value of each 30 second chunk of data was correlated with the bradykinesia and dyskinesia 2 minute scores closest in time to each value. Though we have received 2-minute interval of PKG data we were advised by the manufacturer to use a moving 10 minute average in order to classify patient state. Predefined thresholds supplied by PKG were used on a per patient basis. Bradykinesia scores are negative such that a lower score indicates more bradykinesia. Dyskinesia scores are positive such that more dyskinesia indicated a higher score. Sleep was defined as a bradykinesia score below −80. A patient was scored as “on” with good symptom control by levodopa medication, for each 10 minute segment, if his bradykinesia score was above −26 and his dyskinesia score below 7. Patients were considered “off” if their bradykinesia scores (and/or) dyskinesia scores above 7. Using these guides patient “on” vs “off” state were tailored on an individual basis given each patient’s clinical condition. For example, some patients never had on-time without dyskinesia whereas others did not have any dyskinesia as measured by the PKG. States were defined using a 10 minute window and a two minute step size. A patient was deemed in a certain “state” if at least 3/5 2-minute epochs in a 10 minute window were in agreement with regards to patient state. If this criteria is not met, state is classified as “other” (**Figure 4a**). For each 10-minute window of PKG data the corresponding 30 second PSD’s from the RC+S on the contralateral side were averaged. The use of 30 second PSDs was to avoid transients in the 10 minute windows resulting from small gaps in the data. Thus, we created a moving average with a 10 minute window and 8 minute overlap with average PKG and RC+S scores.

### Supervised motor state detection

Power spectral data from each subject’s individually defined peak in the beta (13-30Hz) frequency in the STN and gamma frequency in cortical contacts was used as input into an linear discriminant analysis (LDA) classifier. Coherence between STN and cortical contacts was also used in these frequency bands. State labels using the PKG wearable data were adjusted according to profile of motor signs (for example not all patients had off-period tremor or on-period dyskinesia). Sleep data (as defined by the PKG wearable) were included in **Figure S1** but are excluded elsewhere. Data from each contact montage, frequency band and measure (power/coherence) were separately fed into an LDA using 5-fold cross validation and the mean AUC (area under the curve) was computed. The same procedure was then repeated for the combination of all features (2 STN beta values, 2 cortical gamma values and 4 coherence measures covering all possible pairings). We used non parametric methods in order to assess the significance of each AUC measurements by repeating the analysis using shuffled labels 5000 times and computing a p-value for each test. Significance was assessed at alpha 0.05 level, corrected for multiple comparisons using the Bonferroni method.

### Unsupervised clustering

In order to perform unsupervised clustering to test whether states are separable without a-priori knowledge of the number of clusters or the location of peaks in the beta frequency the average power in known physiological ranges (delta, theta, alpha, low beta, high beta, beta,low gamma, high gamma) was fed into a clustering detection algorithm^13^. This algorithm does not require specifying the number of clusters (unlike k-means). We used the same temporal averaging as for LDA analysis (a 10 minute window sampled every two minutes). We tested a second clustering approach that relies on capturing known physiological events in clinic and using them as a template to classify at-home data. These states have been well defined in-clinic and correlate with motor impairment in PD but it is unknown if they would be recapitulated in the home environment. We computed the normalized spectral power from a two minute recording in-clinic in which patients were off meds (avoided taking levodopa medication for the past 12 hours) or on meds (after a dose of levodopa medication). We used each state as a template and classified each at-home PSD based on the smallest Euclidian distance to each of the two in-clinic templates.

### Statistical analysis

To assess the statistical significance of in-clinic grouped data we use general estimating equations (GEE) to control for non-independence of our repeated measure test (subject recording site) in the beta and gamma range. We used the same approach to test the significance of at home-grouped data. As input we used the median PSD value from each state classification provided by the PKG watch. We also assessed the significance of each subject’s classification using a non-parametric approach detailed above in *clustering algorithms and state detection.*

### Testing embedded adaptive stimulation

Embedded adaptive stimulation was tested using the protocol of Little et al^14^. that triggered stimulation off of beta bursts^15^. We used the +2-0 STN contact configuration and filtered beta band activity on board the device between 17.58-21.48Hz. Data were sampled at 250Hz with an FFT size of 64 points and three FFT’s were averaged before being input into the onboard linear detector. State change blanking was set at 4 FFT windows to avoid a limit-cycle (self-triggering) in which the stimulation changes are coupled into beta band power estimates. The ramp rate was set at 4.58 mA/sec and the ramp down rate was set at 1.56 mA/sec. The stim rate was set at 130.2 Hz, pulse width at 60us and stim amplitude was set at 0mA below threshold and 2.7 mA above threshold.

The use of “sandwich” mode is important for achieving this, in which strong signal artifact (several orders of magnitude larger than the signal) can be greatly reduced by symmetric bipolar sensing around a monopolar stimulation contact. This does effectively reduce the number and configuration of contacts that can be used for stimulation, when sensing is active.

---

**Figure S1.**
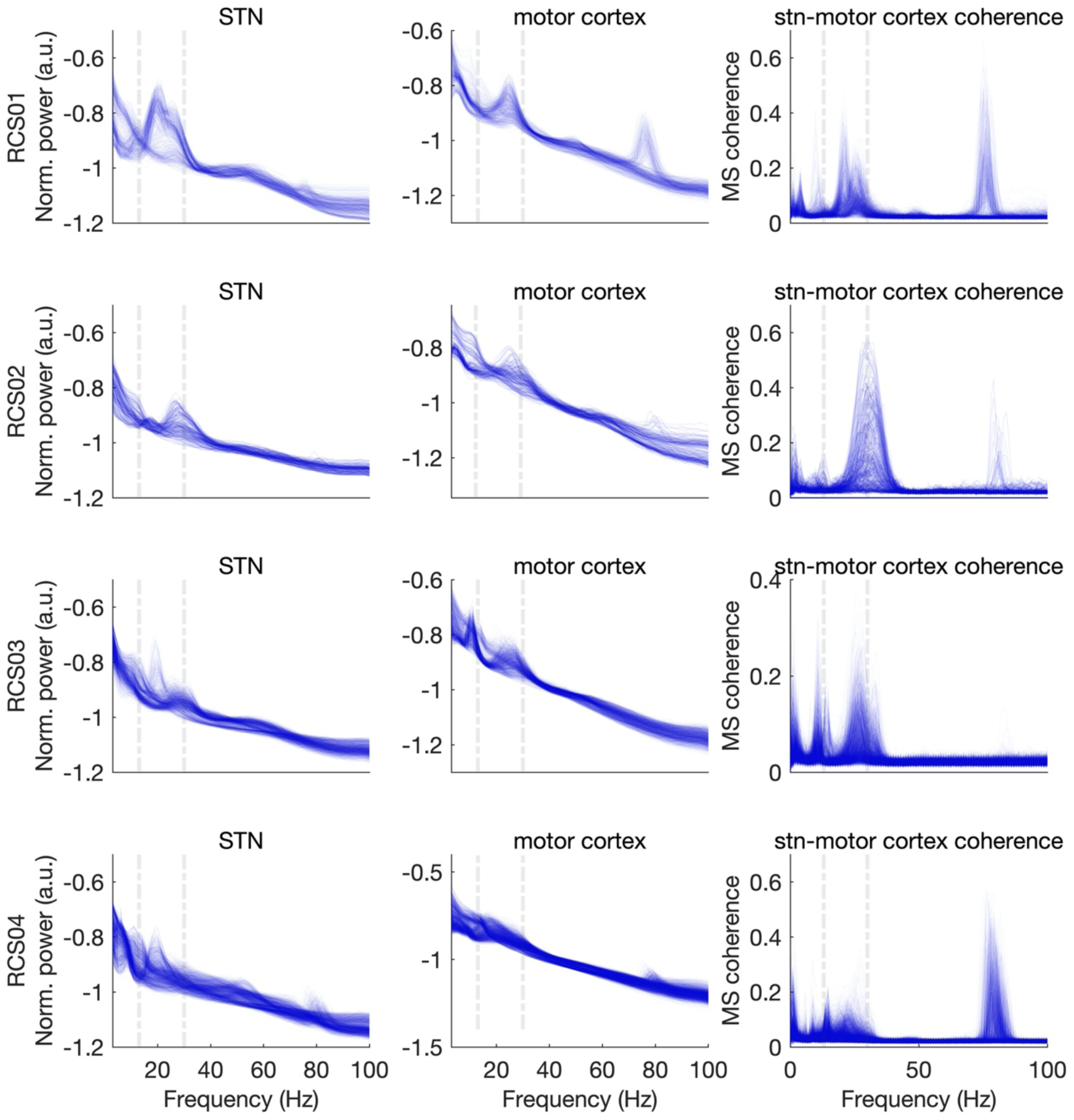
Superimposed STN and motor cortex power spectra (left two columns) and STN-motor cortex coherence (right column) from averaged 10 minute nonoverlapping data segments, showing all data collected during home recordings for all subjects. Both recording channels for each target (0-2 and 1-3 for STN, 8-10 and 9-11 for motor cortex) are represented. Each row shows all data from one study subject. Vertical dotted lines at 13 and 30 Hz demarcate the beta band, for visual clarity.

**Table S1.** Lead locations from fusion of postoperative CT to preoperative MRI.

| subject | AC-PC<br>coordinates<br>of DBS lead<br>tip: left* | AC-PC<br>coordinates<br>of DBS lead<br>tip: right* | Lateral and<br>AP<br>coordinates<br>of lead at<br>dorsal<br>STN:left ** | Lateral and<br>AP<br>coordinates<br>of lead at<br>dorsal<br>STN:right** | Sagittal<br>plane<br>approach<br>angle<br>(degrees<br>from<br>horizontal) | Coronal<br>plane<br>approach<br>angle<br>(degrees<br>from<br>vertical) | Ecog<br>laterality<br>(mm) at<br>central<br>sulcus: left | ECog<br>laterality at<br>central<br>sulcus: right |
| --- | --- | --- | --- | --- | --- | --- | --- | --- |
| RCS01 | -9.8, -3.3, -<br>6.5 | -9.1, -2.5, -<br>6.9 | -10.1, -2.4 | 10.2, -1.1 | Left: 60.7<br>Right:67.1 | Left:-14.6<br>Right:20.0 | 29.0 mm<br>(contact 1) | 24.8 mm<br>(contact 1) |
| RCS02 | -11.8, -3.4,<br>-5.8 | 9.1, -4.0, -<br>5.6 | -11.4, -2.7 | 10.9,-3.2 | Left: 51.4<br>Right: 52.9 | Left: -17<br>Right: 8.7 | 21.1<br>(contact 1) | 28.9 (contact<br>1) |
| RCS03 | -10.2, -3.9,<br>-7.0 | 9.4,-1.4, -<br>6.6 | -10.9, -3.2 | 10.4, -0.2 | Left:70.6<br>Right: 61.6 | Left: 16.6<br>Right:<br>21.6 | 25.4<br>(contact 2) | 29.3<br>(contact 2) |
| RCS04 | -10.5, -3.0,<br>-6.8 | 9.4, -1.1, -<br>7.8 | -11.3, -1.5 | 10.2, 0 | Left: 64.8<br>Right:66.5 | Left:16.7<br>Right:14.4 | 33.1<br>(contact 1) | 22.3 (contact<br>1) |

\*coordinates are lateral, anteroposterior, and vertical with respect to midpoint of line connecting the anterior and posterior commissures
\*\* coordinates are lateral and anteroposterior, in the axial plane 4 mm inferior to the line connecting anterior and posterior commissures

